# The anti-metabolite KAT/3BP has *in vitro* and *in vivo* anti-tumor activity in lymphoma models

**DOI:** 10.1101/2025.04.23.650181

**Authors:** Chiara Tarantelli, Filippo Spriano, Elisa Civanelli, Luca Aresu, Giorgia Risi, Eleonora Cannas, Omar Kayali, Luciano Cascione, Alberto J. Arribas, Anastasios Stathis, Young H. Ko, Francesco Bertoni

## Abstract

Reprogramming of cellular metabolism is a hallmark of cancer, offering therapeutic opportunities to target cancer cell vulnerabilities for therapeutic purposes. 3-Bromopyruvate (3BP), a small alkylating agent, acts as an anti-metabolite to vital substrates in cancer metabolism and exhibits antitumor activity across various cancer types, but the unformulated 3BP can cause high toxicity. This study explores the efficacy of the 3BP clinical derivative KAT/3BP, currently in phase 1 for patients with hepatocellular carcinoma, in lymphoma models. *In vitro*, KAT/3BP demonstrated cytotoxicity across 12 lymphoma cell lines, including diffuse large B-cell lymphoma and mantle cell lymphoma, with a median IC50 of 3.7 μM. KAT/3BP was also effective against lymphoma cell lines with acquired resistance to FDA-approved therapies. *In vivo*, we observed reduced tumor size in a syngeneic mouse model, with the combination of oral and intratumoral administration proving most effective. Additionally, KAT/3BP exhibited synergistic activity when combined with lymphoma therapies, including bendamustine and R-CHOP. These findings underscore the potential of KAT/3BP as a novel therapeutic option for a single agent or a combination of lymphomas.

## Introduction

Reprogramming of cellular metabolism is one of the hallmarks of cancer, and it can provide therapeutic opportunities to exploit cancer cell vulnerabilities (1-3). The 3-Bromopyruvate (3BP) is a small, highly reactive alkylating agent formed by the bromination of pyruvate (4-10). Due to its high structural similarity with pyruvate (Krebs cycle), lactate (Warburg effect), and acetate (lipogenesis), 3BP acts as an anti-metabolite to these vital metabolic substrates for cancer cells. Moreover, since it is a potent alkylating agent, it can modify many proteins, including glycolytic and mitochondrial enzymes. At acidic extracellular pH, 3BP enters cancer cells via monocarboxylic acid-1 (MCT-1) and inhibits glycolysis through hexokinase II (HK-2) covalent modification, with HK-2 inhibition and dissociation from mitochondria, apoptosis-inducing factor (AIF) release, and apoptosis induction (9). Preclinical antitumor activity as a single agent and in combination has been reported in solid tumors, multiple myeloma, and leukemias (4,11-17). In lymphomas, *in vitro* and in vivo antitumor activity has been observed in the Burkitt lymphoma Raji cell line (18,19) and a mouse syngeneic T-cell lymphoma (20,21). In the latter, tumor growth was in vivo reversed, with an increase in the number of circulating CD4+, CD8+, and NK-cells and in the density of tumor-associated macrophages, with reduced local immunosuppression (20). Although the high metabolic activity of cancer cells should allow a preferential effect on neoplastic than healthy cells, unformulated 3BP administrations are associated with severe toxicities, including deaths (22,23). However, improvements have been made in developing novel 3BP formulations based on liposomes, polyethylene glycol (PEG), PEGylated liposomes (stealth liposomes), perillyl alcohol formulations, and others (12,22,24). Thanks to these steps forward, a patient with fibrolamellar hepatocellular carcinoma has been safely treated with a novel formulated 3BP via transcatheter arterial chemoembolization (24). KAT-101 and KAT-201 are two clinical 3BP derivatives formulated for oral or intratumoral (IT) administration, respectively (National Cancer Institute Thesaurus Codes C193479 and C193479), now entering the early clinical evaluation of patients with hepatocellular carcinoma (NCT05603572).

Interestingly, the 3BP-target HK-2 is a metabolic driver in diffuse large B-cell lymphoma (DLBCL) cell lines and patients (25). Moreover, an immunohistochemistry study of 120 clinical specimens showed that the 3BP-transporter MCT1 is expressed in all DLBCL cases (26). Also, considering the limited preclinical data available in lymphomas using 3BP-based therapies, we explored the antitumor activity of the 3BP derivative KAT/3BP in lymphoma models, including cell lines with secondary resistance to FDA-approved agents and a syngeneic mouse model.

## Material and methods

### Cell lines

Lymphoma cell lines were cultured according to the recommended conditions, as previously described (27). All media were supplemented with fetal bovine serum (FBS) (10% or 20%) and penicillin-streptomycin-neomycin (≈5,000 units penicillin, 5 mg streptomycin, and 10 mg neomycin/mL; Sigma-Aldrich, Darmstadt, Germany). Cell line identities were confirmed by short tandem repeat DNA fingerprinting using the Promega GenePrint 10 System kit (B9510). Cells were periodically tested for mycoplasma negativity using the MycoAlert Mycoplasma Detection Kit (Lonza, Visp, Switzerland).

### Compounds

KAT/3BPwas provided by KoDisc overy, LLC. Tazemetostat, ibrutinib, lenalidomide, bendamustine, copanlisib, venetoclax, vorinostat, doxorubicin, vincristine, and prednisolone were purchased from Selleckchem (Houston, TX, USA). Rituximab was purchased from Roche (Basel, Switzerland), and 4-hydroperoxy-cyclophosphamide from Santa Cruz Biotechnology (Heidelberg, Germany).

### *In vitro* cytotoxic activity

Cells were manually seeded in 96-well plates at a concentration of 50,000 cells/mL (10,000 cells in each well). Treatments were performed manually. After 72 hours, cell viability was determined by using 3-(4,5-dimethyl-thiazol-2-yl)-2,5-diphenyltetrazolium bromide, and the reaction stopped after 4 hours with sodium dodecyl sulfate lysis buffer. For R-CHOP treatment, cells were exposed for 72 h to 1 μg/mL CHOP + 100 μg/mL rituximab to five different concentrations in serial dilution 1:10. Rituximab was diluted to clinically recommended serum levels (28) and CHOP represented a mix reflecting the clinical ratios of the drugs (85%, 4-hydroperoxy-cyclophosphamide; 5.5%, doxorubicin; 0.16%, vincristine; 11.1%, prednisolone) (29,30).

For combination studies, cells were exposed (72 hours) to eight increasing concentrations of the two agents, either alone or in combination, followed by an MTT assay. ZIP, HAS, Loewe, and Bliss parameters were calculated for a fixed dose of KAT/3BP, giving anti-proliferative activity between 70 and 10% using SynergyFinder software (31,32).

### Cell cycle and apoptosis assessment

Analysis of apoptosis induction and cell cycle analysis were performed after 24, 48, and 72h of drug exposure at 5μM or DMSO. For apoptosis analysis, cells were stained with Annexin V and propidium iodide, and the percentage of apoptotic cells (Annexin V positive/PI negative and Annexin V positive/PI positive) was assessed. For cell cycle analysis, cells were fixed and permeabilized with ethanol and subsequently stained with propidium iodide. The percentage of cells in sub-G0, G1, S, or G2-M phases was assessed.

### *In vivo* syngeneic mouse models

KAT/3BP was prepared by dissolving 3BP in a buffer system based on sodium phosphate and sodium citrate for oral, IP, and IT deliveries. Mice maintenance and animal experiments were performed under the institutional guidelines established for the Animal Facility at The Institute of Research in Biomedicine (IRB), license national number 30551. BALB/cAnNCrl mice were obtained from Charles River (Calco, Lecco, Italy). Tumors were established by injecting A20 murine lymphoma cells (5 × 10^6^ cells/mouse, 100 μL of PBS) into the left flanks of female BALB/c mice (6-8 weeks of age, approximately 20 gr of body weight). Treatments started once tumor volume reached approximately 60 mm^3^ for pilot treatments and 140 mm^3^ for combination studies, as an average for each group. Tumor volume was measured three times per week using a digital caliper, and animal body weight was measured three times per week throughout the study. The animal status was carefully evaluated during housing and treatments by measuring Cumulative Condition Scores (CCS). Mice were sacrificed once tumor volume reached 2000 mm^3^ and/or when several parameters were scored with a high severity degree (body weight loss, body condition score, physical condition, behavior, hydration, respiration). For IT administration, the endpoint was set at 1500 mm^3^. Tumor samples were fixed in buffered formalin and examined histologically using hematoxylin and eosin staining. A semi-quantitative scoring system, ranging from 0 to 3, was employed to assess the extent of necrosis, corresponding to the percentage of tissue area affected by the necrotic process. The scoring criteria were as follows: score 0 indicated no necrosis present; score 1 represented 1-10% of the tissue involved; score 2 denoted 11-20% of the tissue involved; and score 3 signified approximately 25% of the tissue affected. The assessment was conducted across ten microscopic fields at 40x magnification.

### Statistical analysis

Statistical analyses, including IC50s determination, were conducted using Prism software v10.2.3 (GraphPad Software, La Jolla, CA, USA). Statistical significance was determined by a two-tailed unpaired Student’s t-test or as described in the figure legends. A P value < 0.05 was considered statistically significant.

## Results

### KAT/3BP as a single agent is cytotoxic in lymphoma cell lines

Twelve lymphoma cell lines derived from activated B-cell-like diffuse large B-cell lymphoma (ABC DLBCL) (n = 4; TMD8, RCK8, U2932, SUDHL2), germinal center B-cell like (GCB) DLBCL (n = 4; OCILY19, WSU-DLCL2, DOHH2, TOLEDO), and mantle cell lymphoma (MCL) (n = 4; MINO, REC1, GRANTA519, Z138), were exposed to increasing concentrations of the 3BP clinical derivative KAT/3BP for 72 hours (Figure 1). The median IC50 across all the cell lines was 3.7 μM, with no differences based on the histotypes.

**Figure 1.**
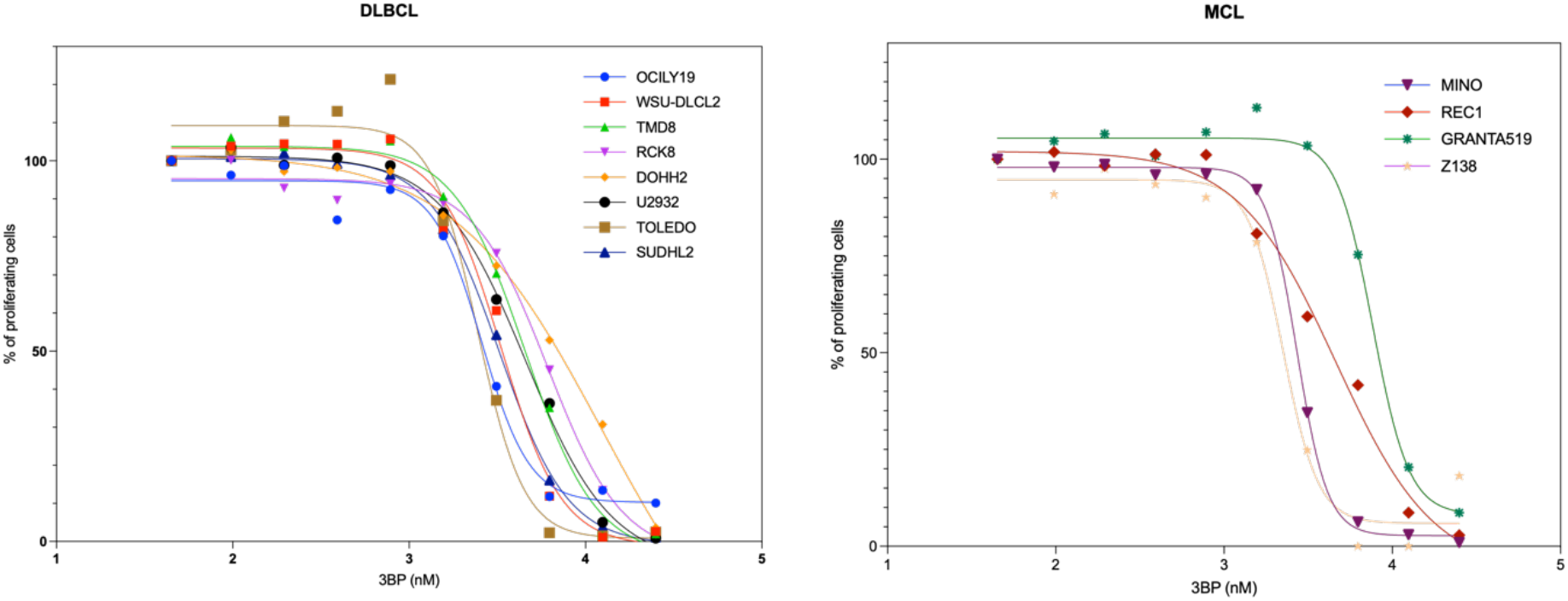
Antiproliferative effect of KAT/3BP in DLBCL and MCL subtypes of lymphoma. The dose-response curve in four mantle cell lymphoma (right panel) and eight diffuse large B cell lymphoma cell lines (right panel) treated with KAT/3BP treated with increasing KAT/3BP compound concentrations. MTT assay was performed to evaluate the anti-tumoral activity of the drug.

Cell cycle analyses in one DLBCL (Toledo) and one MCL (Z138) cell line exposed to KAT/3BP at 5 μM or DMSO as a control for 24, 48, and 72 hours showed a time-dependent increase in the percentage of cells in sub-G0 (Supplementary Figure 1A). Thus, an Annexin V test showed an apoptosis induction in both cell lines already at 24 hours of exposure to KAT/3BP at 5 μM (Supplementary Figure 1B).

### KAT/3BP, as a single agent, exerts anti-lymphoma activity in models of secondary resistance to FDA-approved agents

Based on the activity observed in DLBCL and MCL cell lines, we also tested the compound in two marginal zone lymphoma (MZL) cell lines (Karpas1718 and VL51) and their derivatives with acquired resistance to PI3K inhibitors, BTK inhibitors, and BCL2 inhibitors (33-36). KAT/3BP showed a dose-dependent anti-proliferative activity in the two MZL cell lines and the resistant cells (Figure 2).

**Figure 2.**
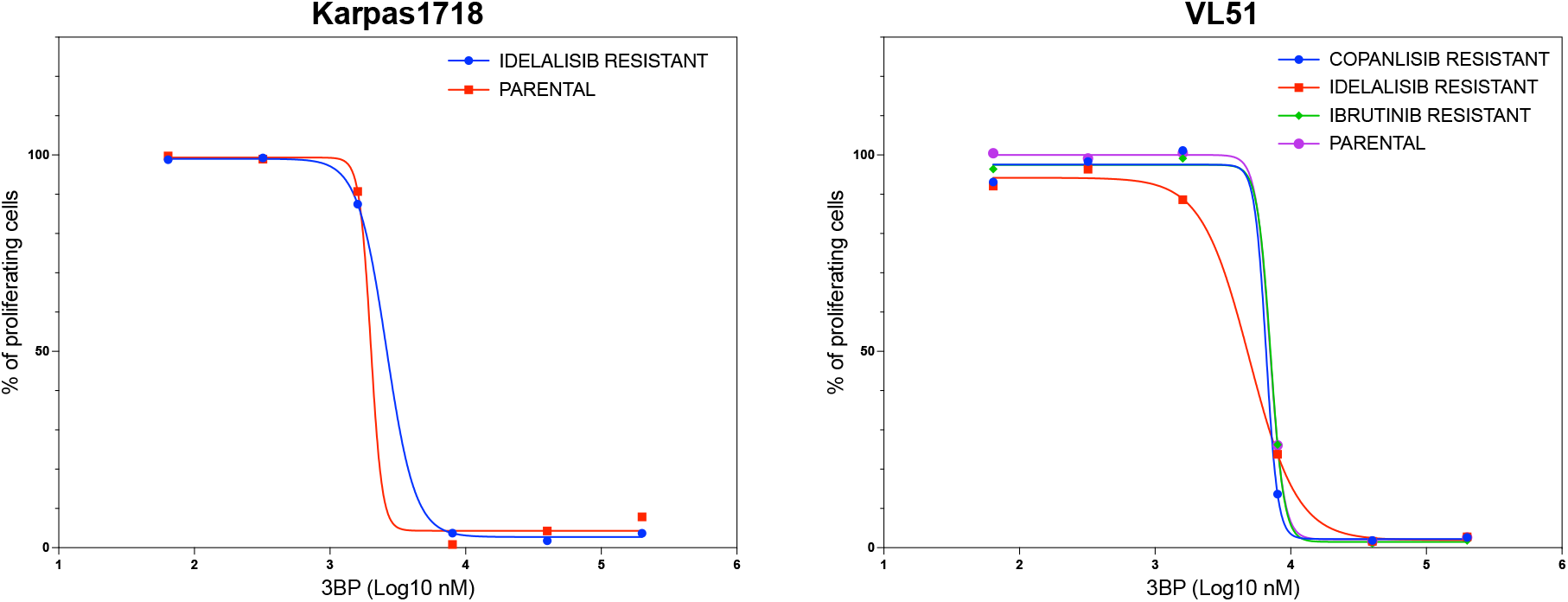
Antiproliferative effect of KAT/3BP in MZL models of secondary resistance to FDA approved agents. Marginal zone lymphoma cell lines (parental and resistant to idelalisib, ibrutinib, and copanlisib) were treated with increasing KAT/3BP compound concentrations. MTT assay was performed to evaluate the anti-tumoral activity of the drug.

### KAT/3BP has *in vivo* antitumor activity in a syngeneic mouse model

We then validated the *in vitro* results using a murine syngeneic model (A20 lymphoma cells, BALB/c mice). First, we confirmed that KAT/3BP was also *in vitro* active in the A20 lymphoma cells (Supplementary Figure 2).

Mice were treated with KAT/3BP as oral, IP, or IT delivery route or with a buffer as a control for 28 days. All mice in the control groups survived up to day 26 for the IP administration group and day 19 for the PO and IT groups.

Treatments with oral or IT KAT/3BP administration determined reduced tumor size compared to the control groups (Figure 3A-B, Supplementary Figure 3). This effect was further evidenced by the slope values extrapolated from a linear regression model (Table 1). Treatments with oral administration at 10 mg/kg dose led to a total tumor reduction in three mice out of five; two were still alive at day 92, and one was sacrificed on day 45. The latter had a relapse in the upper part of the thorax. One mouse in the IT low group and another in the PO + IT low had to be sacrificed due to tumor growth, which occurred much later than what was seen in the control group.

**Table 1.** Estimated slopes for each group under a single administration route of 3BP in an *in vivo* syngeneic model. Calculated slopes following a linear regression model are shown in the table.

|  | CTRL PO | PO low | PO high | CTRL IT | IT low | IT high | PO + IT low | PO + IT high |
| --- | --- | --- | --- | --- | --- | --- | --- | --- |
| Best-fit values |  |  |  |  |  |  |  |  |
| Slope | 101.8 | 67.84 | -2.610 | 68.24 | 18.71 | 39.26 | 13.95 | 22.39 |

**Figure 3.**
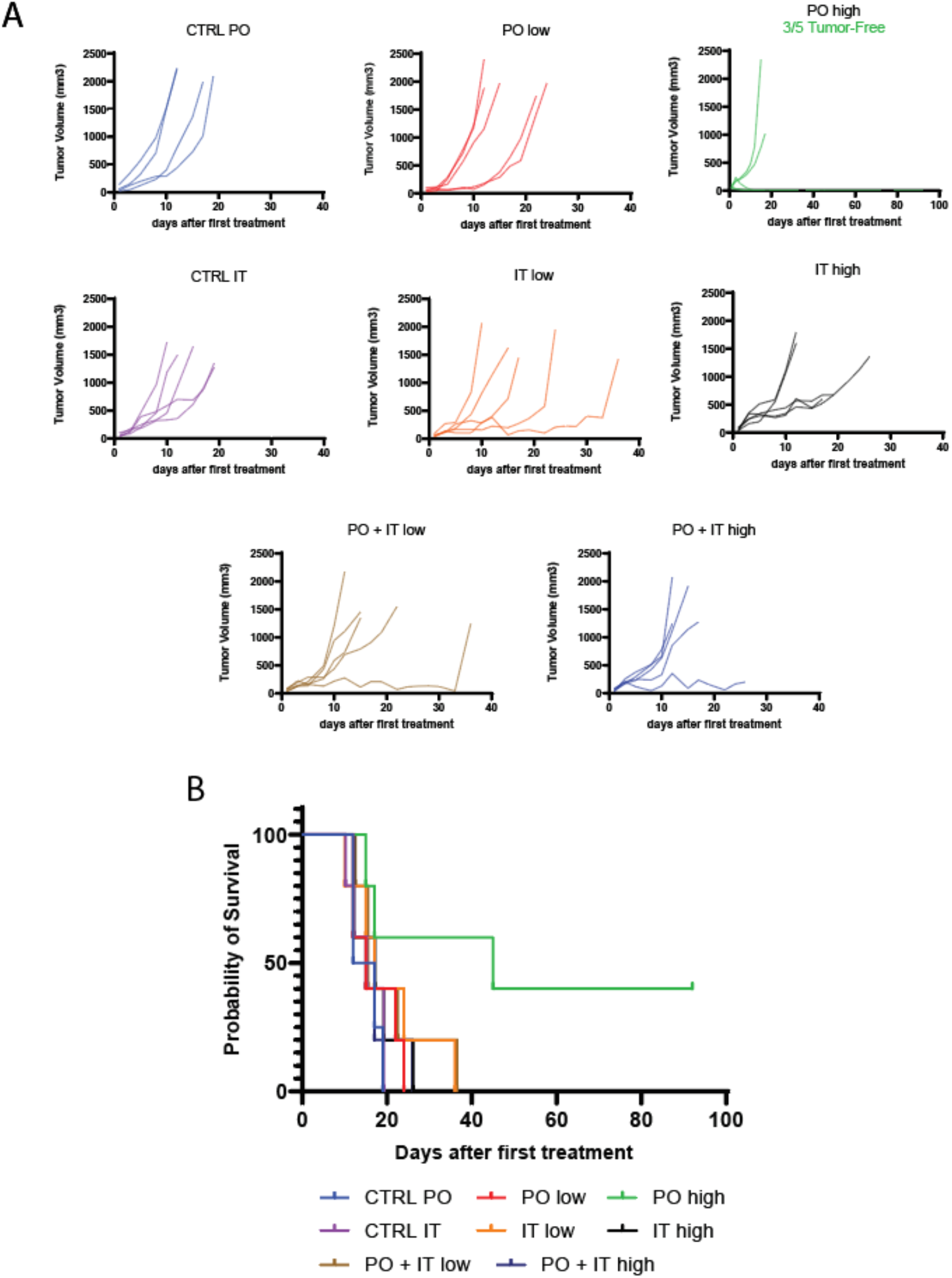
Assessment of KAT/3BP anti-lymphoma activity in *in vivo* syngeneic model. BALB/c mice were subcutaneously injected with murine lymphoma cell line A20. Mice were treated with vehicle (SFB) by oral or intratumoral (IT) injection, with 2.5 mg/kg and 10 mg/kg per os (low and high PO, respectively), with 0.5 mM and 2 mM by IT (low and high IT, respectively), with the combination of 2.5 mg/kg PO plus 0.5 mM IT, or with the combination of 10 mg/kg PO and 2 mM IT (PO + IT low and PO + IT high, respectively). All groups were composed by five mice in total, and the end of the experiment was set at day 92 for mice with complete tumor regression. (A) Graphs showing tumor volume in mm^3^ for each animal in each group. (B) Survival for each group.

Necrosis was observed in both the periphery and center of the tumor, exhibiting a multifocal distribution. The PO + IT high, PO high, and IT high groups (one sample each) were characterized by necrosis. However, a score of 3, indicating the highest level of necrosis, was only observed in both the IT high and PO + IT high groups (Supplementary Figure 4).

IP administration was not tolerated at a high dose of 10 mg/kg, with 10-15% body weight loss after three days of treatment, dehydration, and a cumulative condition score at the maximum limit (Supplementary Figure 5A-C). IP low dose group showed a 5% body weight loss after three days and recovered after eating soft food in the cage. Treatment was not effective in tumor reduction.

We then evaluated the combination of oral and IT KAT/3BP. Groups of eight mice each were treated orally and intratumorally for 28 days with the vehicle, oral KAT/3BP administered at a high dosage (10 mg/kg), IT KAT/3BP at a low dosage (0.5 mM), and the combination of the two. Treatments with oral plus IT KAT/3BP decreased tumor volume growth compared to the other groups, with complete tumor reduction in one mouse out of nine of the combination groups (Figure 4A, Supplementary Figure 6A). Based on the slope calculated with linear regression mode, a higher tumor reduction than the control vehicle was observed in all the treatment groups, especially in the combination group (Table 2).

**Figure 4.**
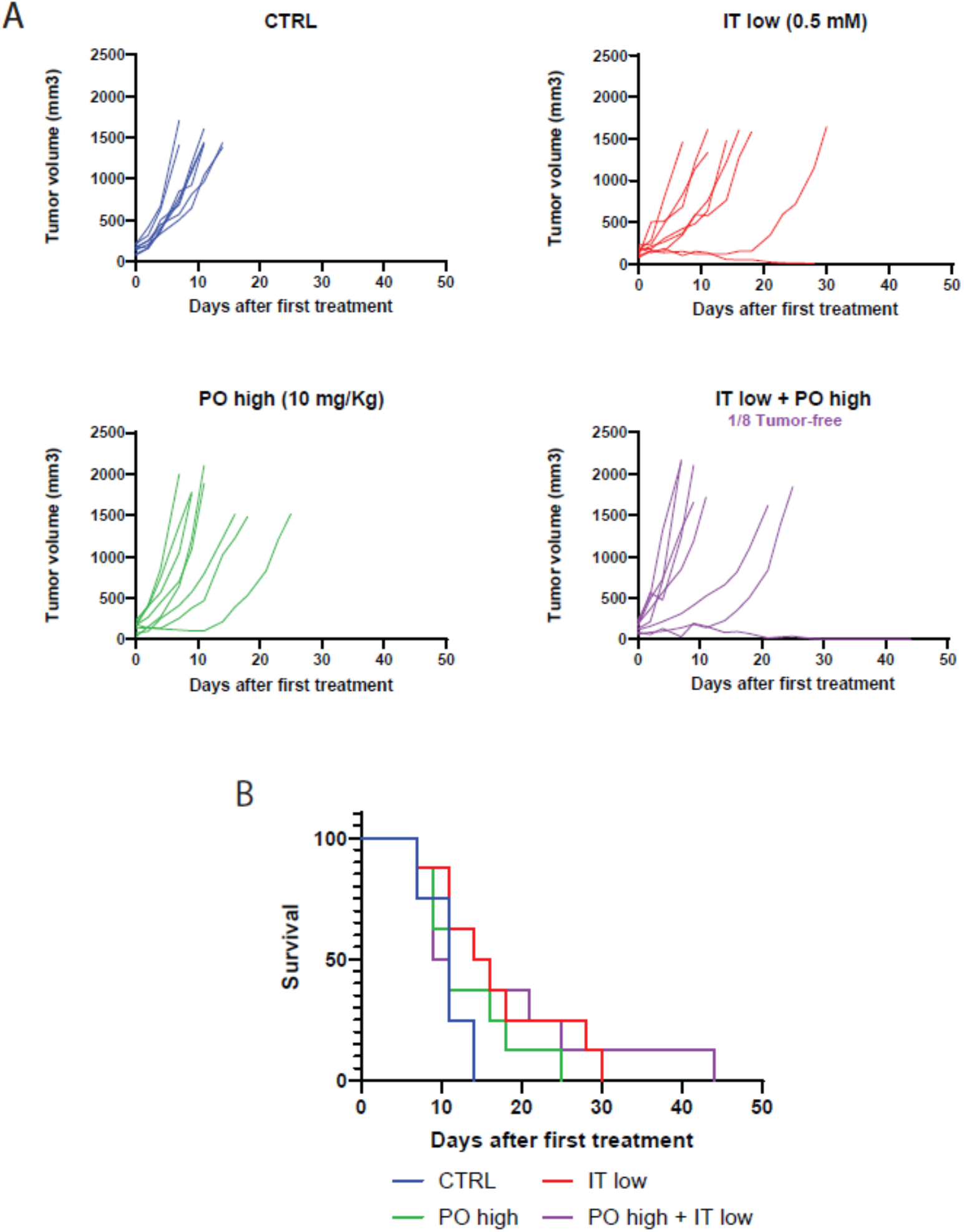
Assessment of KAT/3BP anti-lymphoma activity in PO and IT administration as single and in combination. BALB/c mice were subcutaneously injected with murine lymphoma cell line A20. Mice were treated with vehicle (SFB) by oral or intratumoral (IT) injection, with 10 mg/kg per os (PO high), with 0.5 mM by IT (IT low), with the combination of PO high plus IT low, or with the combination of the two (PO high + IT low). All groups were composed by 9 mice in total, and the end of the experiment was set at day 92 for mice with complete tumor regression. (A) Graphs showing tumor volume in mm^3^ for each animal in each group. (B) Survival for each group. End of the experiment: D44.

**Table 2.** Slope values are extrapolated by a linear regression model in PO and IT KAT/3BP administration as single and combination routes. Calculated slopes following a linear regression model are shown in the table.

|  | CTRL PO | PO low | PO high | CTRL IT |
| --- | --- | --- | --- | --- |
| Best-fit values |  |  |  |  |
| Slope | 101.8 | 67.84 | -2.610 | 68.24 |

The survival graph shows that all the treated groups of mice, regardless of the administration route, have better survival than the vehicle group (Figure 4B). In the combination group, one mouse survived up to day 44. The experiment was stopped due to license limitations, which prevented single-housed animals from being kept. The animal was visually inspected for any presence of a tumor, and the absence of a cancer mass was confirmed. One mouse in the IT group developed a second tumor and was sacrificed on day 28 when the maximum volume was unacceptable.

Tumor ulcerations were detected in two mice in the vehicle group (PO high + IT low administration), four in the IT group, one in the oral group, and two mice in the combination. Mice’s weight was monitored, and the drug did not induce weight loss of more than 5% (Supplementary Figure 6B).

### KAT/3BP-based combinations are active in lymphoma cell lines

Based on the demonstration of *in vitro* and *in vivo* single-agent activity, we explored possible KAT/3BP-based combinations. We tested FDA-approved agents or molecules targeting critical pathways in lymphoma. Cell lines were exposed for 72 hours to a fixed dose of KAT/3BP, increasing concentrations of the second drug as single agents and combined. The latter included the EZH2 inhibitor tazemetostat, BTK inhibitor ibrutinib, the cereblon E3 ligase modulator lenalidomide, the DNA binding agent bendamustine, the PI3K α/δ inhibitor copanlisib, the BCL2 inhibitor venetoclax, the HDAC inhibitor vorinostat, and the immuno-chemotherapy R-CHOP. Combinations were tested in cell lines derived from GCB (TOLEDO, WSU-DLCL2) and ABC DLBCL (TMD8, U2932). Ibrutinib and lenalidomide were tested only in cell lines derived from ABC DLBCL, while tazemetostat in GCB DLBCL is the subtype in which the drugs have reported clinical activity. Table 3 represents the combination results. For each combination, we have calculated four different synergism parameters: HSA, Bliss, Lowe, and ZIP. Overall, KAT/3BP showed benefits in combination with all the tested compounds. More in detail, we observed synergism in at least one of the four parameters, in combination with R-CHOP in all four DLBCL cell lines. Bendamustine was synergistic in all except TOLEDO. Tazemetostat was synergistic in the TOLEDO. Lenalidomide, ibrutinib, and venetoclax were synergistic in TMD8.

**Table 3.**
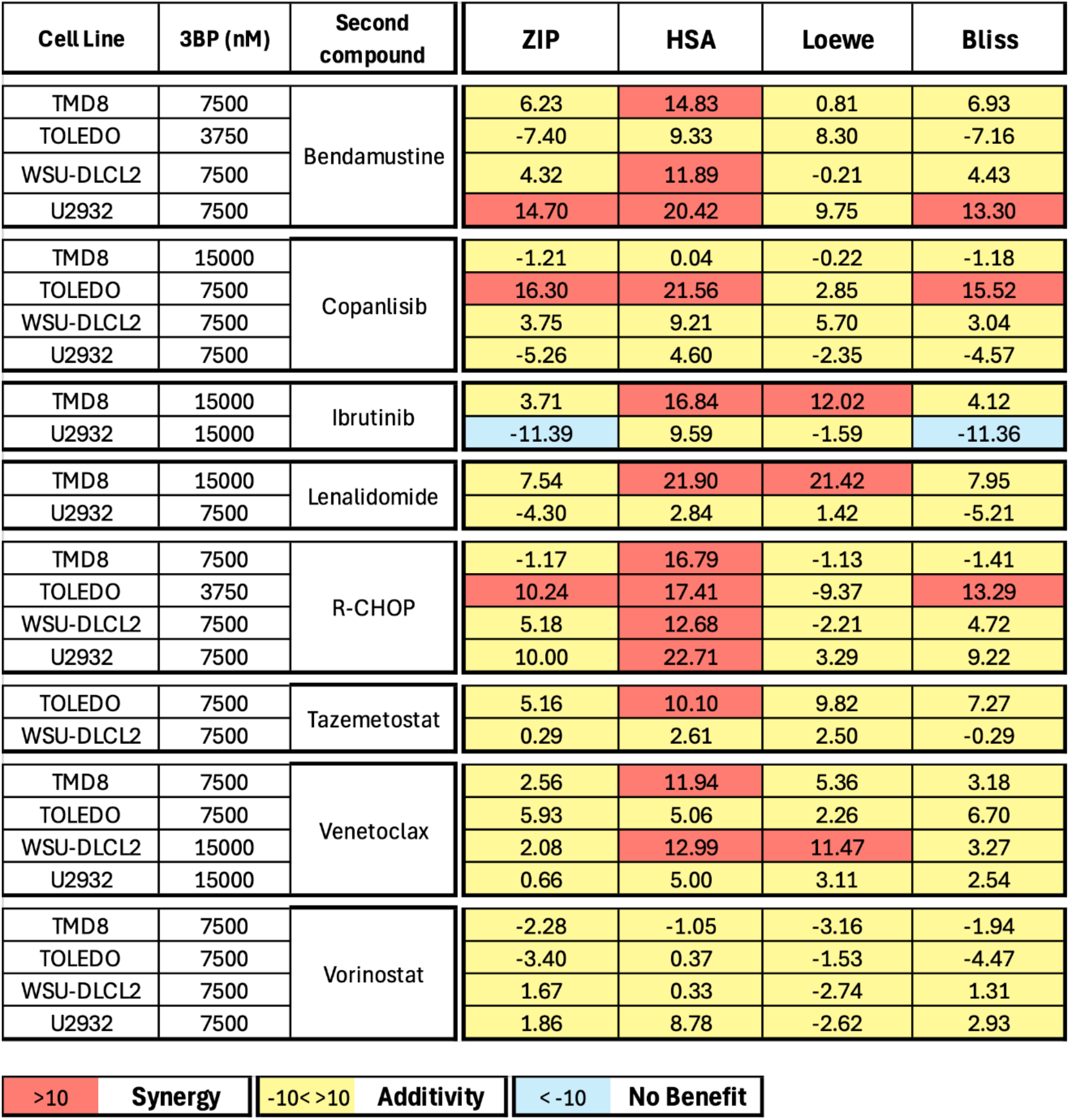
KAT/3BP-containing combinations. The figure shows ZIP, HSA, Loewe, and Bliss combination scores obtained by combining KAT/3BP at a fixed dose with a second compound at increasing doses. Values below -10 indicate no benefit from combining the two drugs; values between -10 and 10 indicate additivity, while values above 10 indicate synergism.

| Cell Line | 3BP (nM) | Second compound | ZIP | HSA | Loewe | Bliss |
| --- | --- | --- | --- | --- | --- | --- |
| TMD8 | 7500 | Bendamustine | 6.23 | 14.83 | 0.81 | 6.93 |
| TOLEDO | 3750 |  | -7.40 | 9.33 | 8.30 | -7.16 |
| WSU-DLCL2 | 7500 |  | 4.32 | 11.89 | -0.21 | 4.43 |
| U2932 | 7500 |  | 14.70 | 20.42 | 9.75 | 13.30 |
| TMD8 | 15000 | Copanlisib | -1.21 | 0.04 | -0.22 | -1.18 |
| TOLEDO | 7500 |  | 16.30 | 21.56 | 2.85 | 15.52 |
| WSU-DLCL2 | 7500 |  | 3.75 | 9.21 | 5.70 | 3.04 |
| U2932 | 7500 |  | -5.26 | 4.60 | -2.35 | -4.57 |
| TMD8 | 15000 | Ibrutinib | 3.71 | 16.84 | 12.02 | 4.12 |
| U2932 | 15000 |  | -11.39 | 9.59 | -1.59 | -11.36 |
| TMD8 | 15000 | Lenalidomide | 7.54 | 21.90 | 21.42 | 7.95 |
| U2932 | 7500 |  | -4.30 | 2.84 | 1.42 | -5.21 |
| TMD8 | 7500 | R-CHOP | -1.17 | 16.79 | -1.13 | -1.41 |
| TOLEDO | 3750 |  | 10.24 | 17.41 | -9.37 | 13.29 |
| WSU-DLCL2 | 7500 |  | 5.18 | 12.68 | -2.21 | 4.72 |
| U2932 | 7500 |  | 10.00 | 22.71 | 3.29 | 9.22 |
| TOLEDO | 7500 | Tazemetostat | 5.16 | 10.10 | 9.82 | 7.27 |
| WSU-DLCL2 | 7500 |  | 0.29 | 2.61 | 2.50 | -0.29 |
| TMD8 | 7500 | Venetoclax | 2.56 | 11.94 | 5.36 | 3.18 |
| TOLEDO | 7500 |  | 5.93 | 5.06 | 2.26 | 6.70 |
| WSU-DLCL2 | 15000 |  | 2.08 | 12.99 | 11.47 | 3.27 |
| U2932 | 15000 |  | 0.66 | 5.00 | 3.11 | 2.54 |
| TMD8 | 7500 | Vorinostat | -2.28 | -1.05 | -3.16 | -1.94 |
| TOLEDO | 7500 |  | -3.40 | 0.37 | -1.53 | -4.47 |
| WSU-DLCL2 | 7500 |  | 1.67 | 0.33 | -2.74 | 1.31 |
| U2932 | 7500 |  | 1.86 | 8.78 | -2.62 | 2.93 |

|  |  |  |  |  |  |
| --- | --- | --- | --- | --- | --- |
| >10 | **Synergy** | -10<>10 | **Additivity** | <-10 | **No Benefit** |

## Discussion

Here, we demonstrated that KAT/3BP has *in vitro* and *in vivo* cytotoxic activity in lymphomas as a single agent, maintained in models with secondary resistance to BTK, BCL2, and PI3K inhibitors. We also showed that adding KAT/3BP to other compounds improves anti-lymphoma effects.

The *in vitro* anti-lymphoma activity of KAT/3BP was seen in the low micromolar range in cell lines derived from three different types of aggressive B cell lymphomas, for which still too many patients need therapies: ABC DLBCL, GCB DLBCL, and MCL (37-39). Indeed, among the models sensitive to KAT/3BP, there were also cell lines that have low sensitivity to the CD19 targeting antibody-drug conjugate loncastuximab tesirine and R-CHOP (U2932, SU-DHL-2) (40)or to CD79B targeting antibody-drug conjugate polatuzumab vedotin (RCK8, SU-DHL-2) (41).

We also observed *in vitro* activity in two cell lines derived from MZL, another lymphoma subtype, and, more importantly, in derivative cell lines that had acquired secondary resistance to BTK inhibitors, PI3K inhibitors, and BCL2 inhibitors. Others have also reported apoptosis in 3BP-treated colorectal cancer models with primary or secondary resistance to EGFR inhibitors (17). The efficacy observed in resistant cell lines further emphasizes KAT/3BP’s potential to overcome resistance mechanisms, a significant challenge in current lymphoma treatments.

The *in vitro* activity was *in vivo* confirmed using a lymphoma syngeneic mouse model. It is worth mentioning that the animals were not fasted before KAT/3BP treatment, although fasting conditions might be more beneficial for the treatment. Oral administration of KAT/3BP at 10 mg/kg effectively reduced the tumor masses, with three mice remaining tumor-free up to day 45 and two remaining tumor-free up to day 92. Furthermore, adding intratumoral KAT/3BP enhanced tumor regression, suggesting that an oral and local drug delivery administration approach could maximize therapeutic outcomes. Conversely, the intraperitoneal administration of KAT/3BP at the dose of 10 mg/kg was acutely toxic, causing body weight loss within only three days from administration, necessitating the sacrifice of the affected animals in this group.

The observed anti-tumor activity of KAT/3BP was primarily cytotoxic, as highlighted by both the *in vitro* apoptosis induction and *in vivo* necrosis in some responding tumors. Similar findings have been reported in various additional tumor models, all pointing towards necrosis as one of the main effects in 3BP-treated cancers (42-44), although the administration route itself might have contributed to the outcome in some mice.

Finally, we demonstrated the benefit of adding KAT/3BP to established anti-lymphoma therapies, especially chemo-based approaches. Indeed, while the anti-tumor effect was improved in combining KAT/3BP with bendamustine or the in vitro version of the R-CHOP regimen, only selected models and targeted agents benefited from adding KAT/3BP.

In conclusion, our data showed that KAT/3BP has *in vitro* and *in vivo* activity in lymphoma models. Moreover, KAT/3BP maintained its activity in models of primary resistance to R-CHOP and antibody-drug conjugates or of secondary resistance to BTK and PI3K inhibitors, and its addition was beneficial when added to standard therapies, mainly chemotherapies. The data support extending the early clinical evaluation, currently running for hepatocellular carcinoma patients, to the lymphoma setting.

## Funding

This work was partially supported by institutional research funds from NewG Lab Pharma, Inc. and KoDiscovery, LLC, and grants from the Swiss National Science Foundation (SNSF 31003A_163232/1) and Swiss Cancer Research (KFS-4727-02-2019).

## Author Contributions

CT, FS: performed experiments, data mining, interpreted data, and co-wrote the manuscript.

AJA: performed experiments and interpreted data.

ECi, GR, ECa, GR: performed experiments. LC: provided advice.

LA: performed experiments, interpreted data, and provided advice.

YHK: co-designed the study, provided reagents, and supervised the study.

FB: co-designed the study, performed data mining, interpreted data, supervised the study, and co-wrote the manuscript.

All authors reviewed and accepted the final version of the manuscript.

## Conflict of interest

CT: travel grant from iOnctura. AJA: travel grant from Astra Zeneca, consultant fee for PentixaPharm. LC: institutional research funds from Orion. AS: institutional research funds for clinical trials: from Abbvie, ADC Therapeutics, Amgen, Astra Zeneca, Bayer, BMS, Cellestia, Debiopharm, Incyte, Loxo Oncology, Merck MSD, Novartis, Pfizer, Philogen, Prelude Therapeutics, Roche; Consultant/expert testimony/advisory board: (institutional): Debiopharm, Janssen, AstraZeneca, Incyte, Eli Lilly, Novartis, Roche, Loxo Oncology; (personal): Incyte Travel grant: Incyte; Astra Zeneca. YHK: president of KoDiscovery, LLC. FB: institutional research funds from ADC Therapeutics, Bayer AG, BeiGene, Floratek Pharma, Helsinn, HTG Molecular Diagnostics, Ideogen AG, Idorsia Pharmaceuticals Ltd., Immagene, ImmunoGen, Menarini Ricerche, Nordic Nanovector ASA, Oncternal Therapeutics, Spexis AG; consultancy fee from BIMINI Biotech, Floratek Pharma, Helsinn, Immagene, Menarini, Vrise Therapeutics; advisory board fees to institution from Novartis; expert statements provided to HTG Molecular Diagnostics; travel grants from Amgen, Astra Zeneca, iOnctura. The other Authors have nothing to disclose.

