## Supplementary tables and figures for "The anti-metabolite KAT/3BP has *in vitro* and *in vivo* anti-tumor activity in lymphoma models"

### Supplementary figures and tables

**Supplementary Figure 1. The activity of KAT/3BP on lymphoma cell lines, one diffuse large B cell lymphoma (TOLEDO), and one mantle cell lymphoma (Z138).** (A) Cell cycle distribution by Propidium Iodide (PI) staining and (B) apoptosis induction by Annexin V/PI FACS acquisition were assessed after 24, 48, and 72 hours of 5 mM drug concentration. Average of two independent experiments.

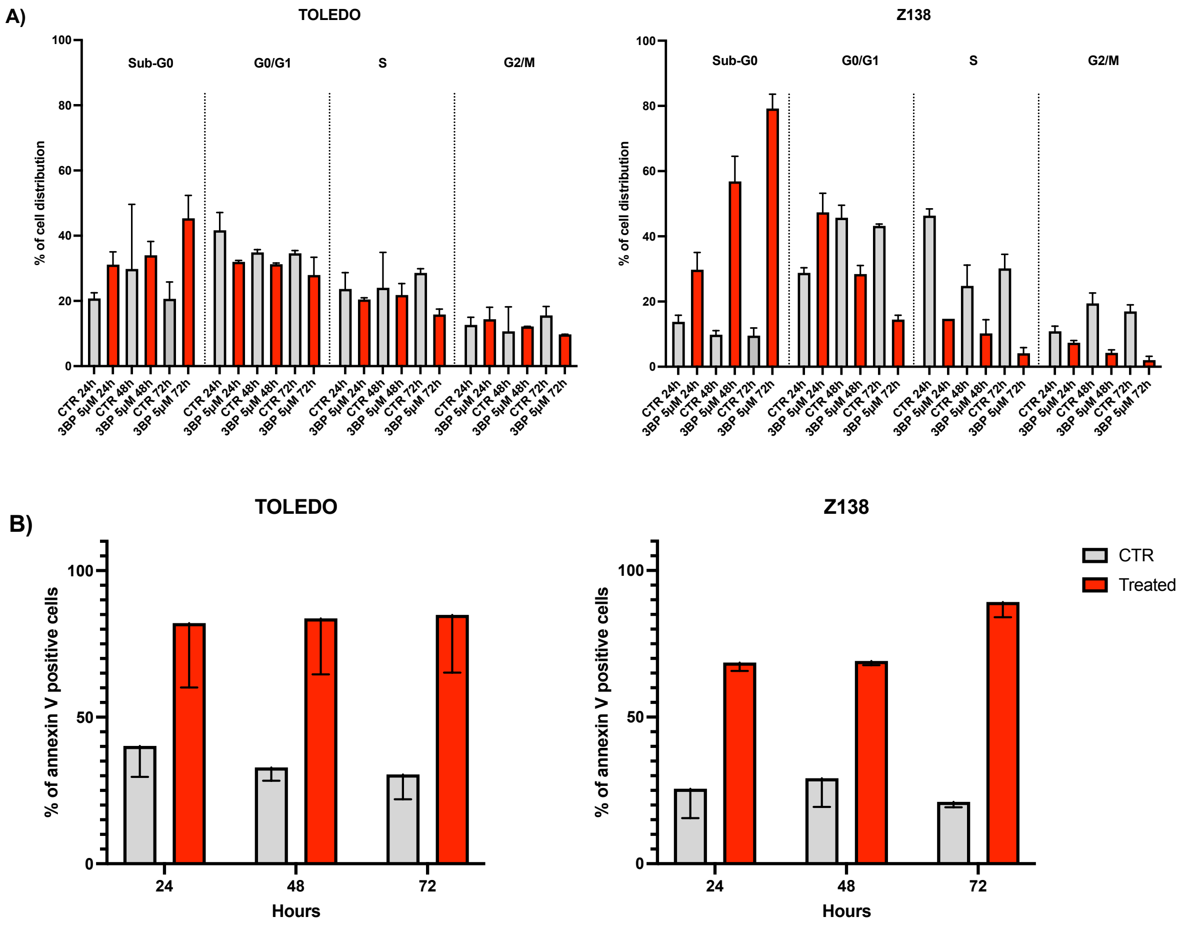

**Supplementary Figure 2. Antiproliferative effect of 3BP in A20 murine cell lymphoma mode.** MTT assay was performed to evaluate the anti-tumoral activity of the drug after 72h of treatment.

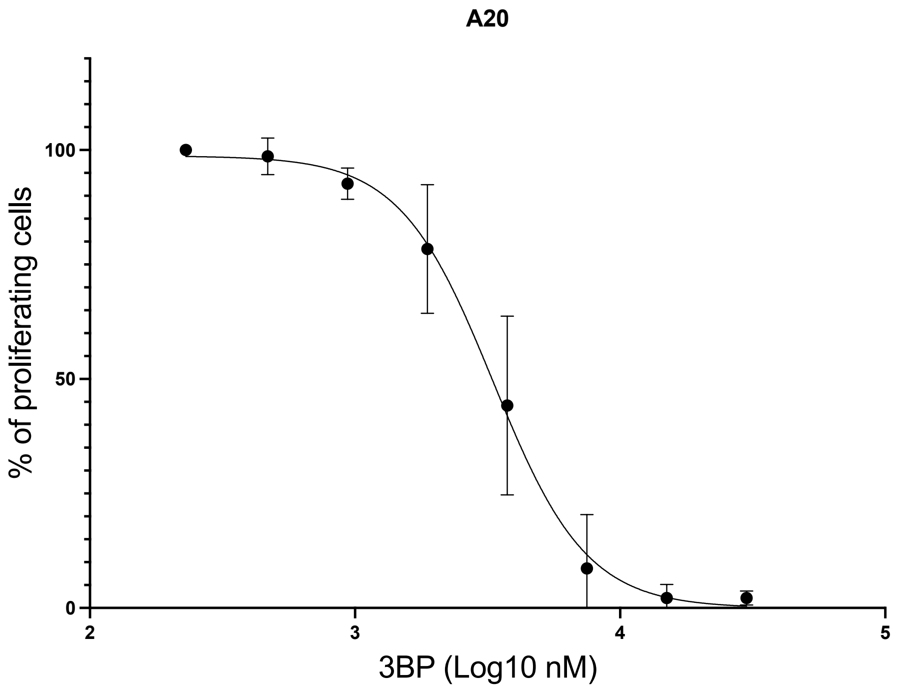

**Supplementary Figure 3. Assessment of KAT/3BP anti-lymphoma activity in *in vivo* syngeneic model.** A) Tumor volume mean with positive and negative SEM for each group. B) Body weight mean with standard deviation for each group. Figure up to the end of the experiment (day 92).

A

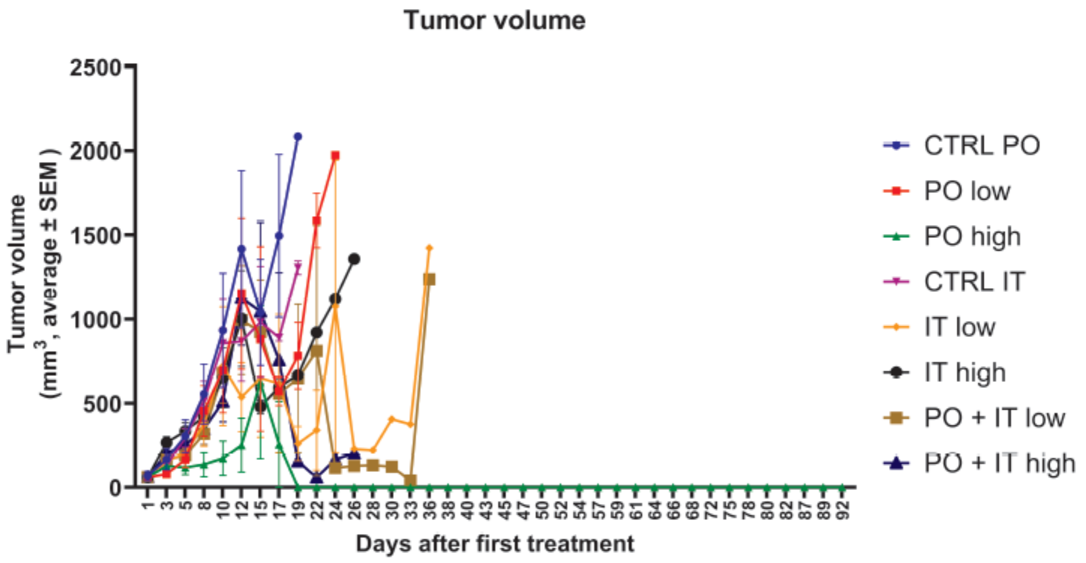


B

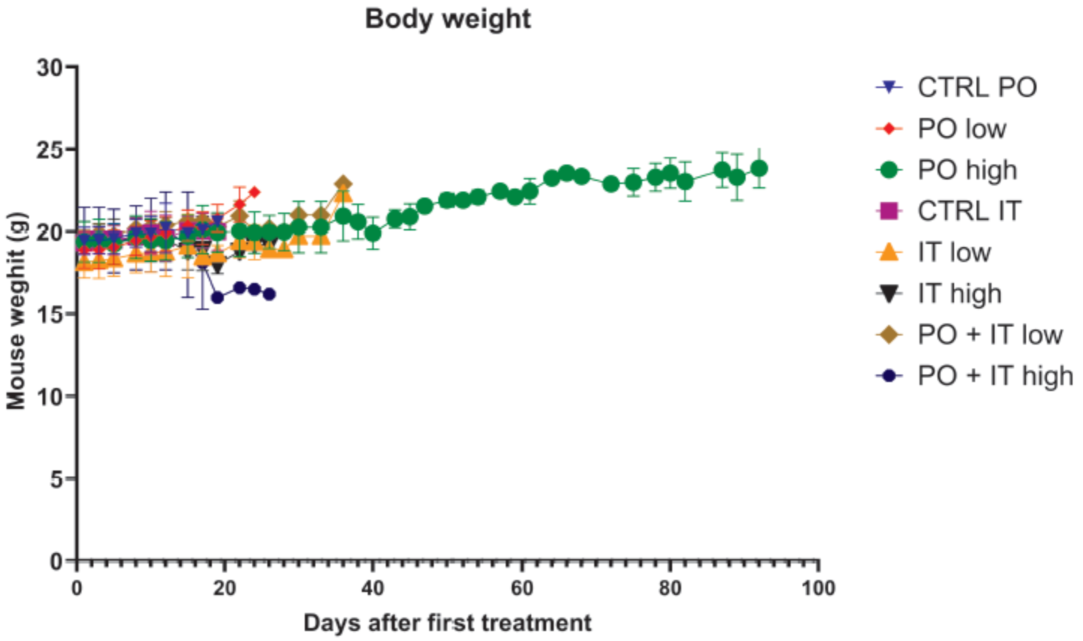

**Supplementary Figure 4. Histological representation of tumor necrosis scores**. A) Score 0: no necrosis present. The tumor tissue shows intact cellular architecture without evidence of cell death or tissue breakdown. B) Score 1: The focal area of cellular degradation and tissue disruption are visible, affecting up to 10% of the examined field. C) Score 2: more extensive areas of cell death and tissue disorganization are apparent. D) Score 3: widespread necrotic regions are evident, characterized by extensive cellular debris, loss of nuclear detail, and disrupted tissue architecture.

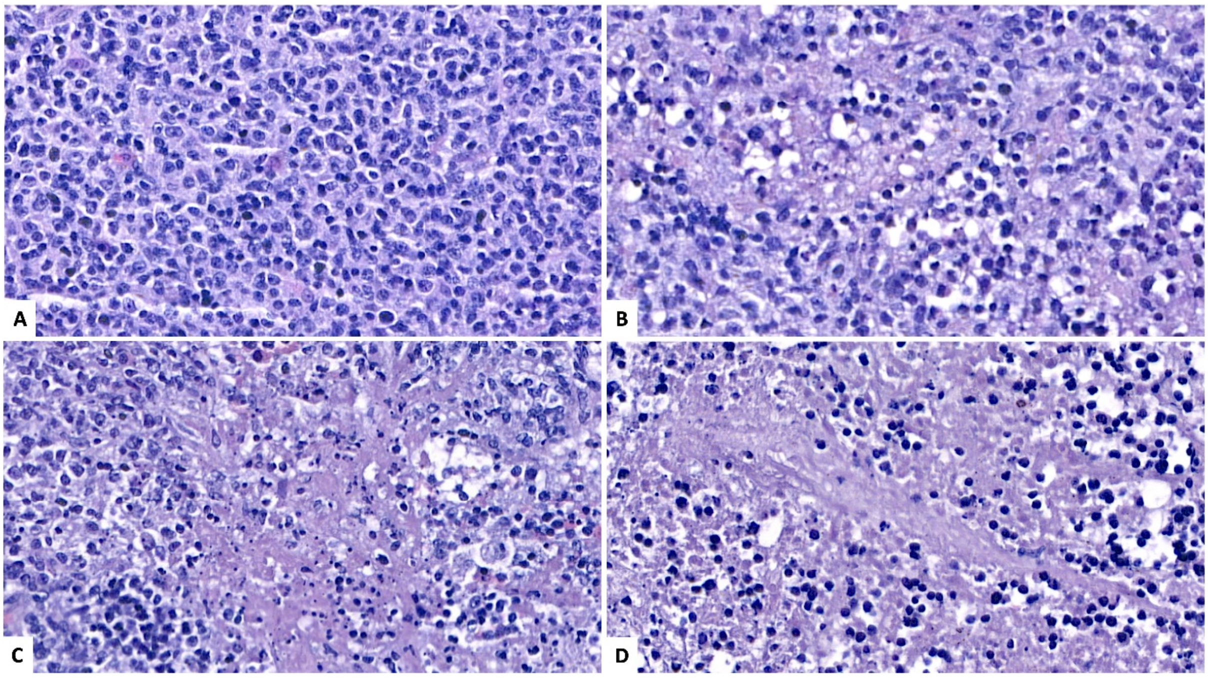

**Supplementary Figure 5. Activity of KAT/3BP administered by intraperitoneal injection in a syngeneic model.** BALB/c mice were subcutaneously injected with murine lymphoma cell line A20. Mice were treated with vehicle (SFB) by intraperitoneal (IP) injection, with 2.5 mg/kg and 10 mg/kg (low and high IP, respectively). A) Graphs showing tumor volume in $mm^3$ for each animal in each group. B) Probability of survival for each group. C) Body weight mean with standard deviation for each group. Figure up to the end of the experiment (day 26).

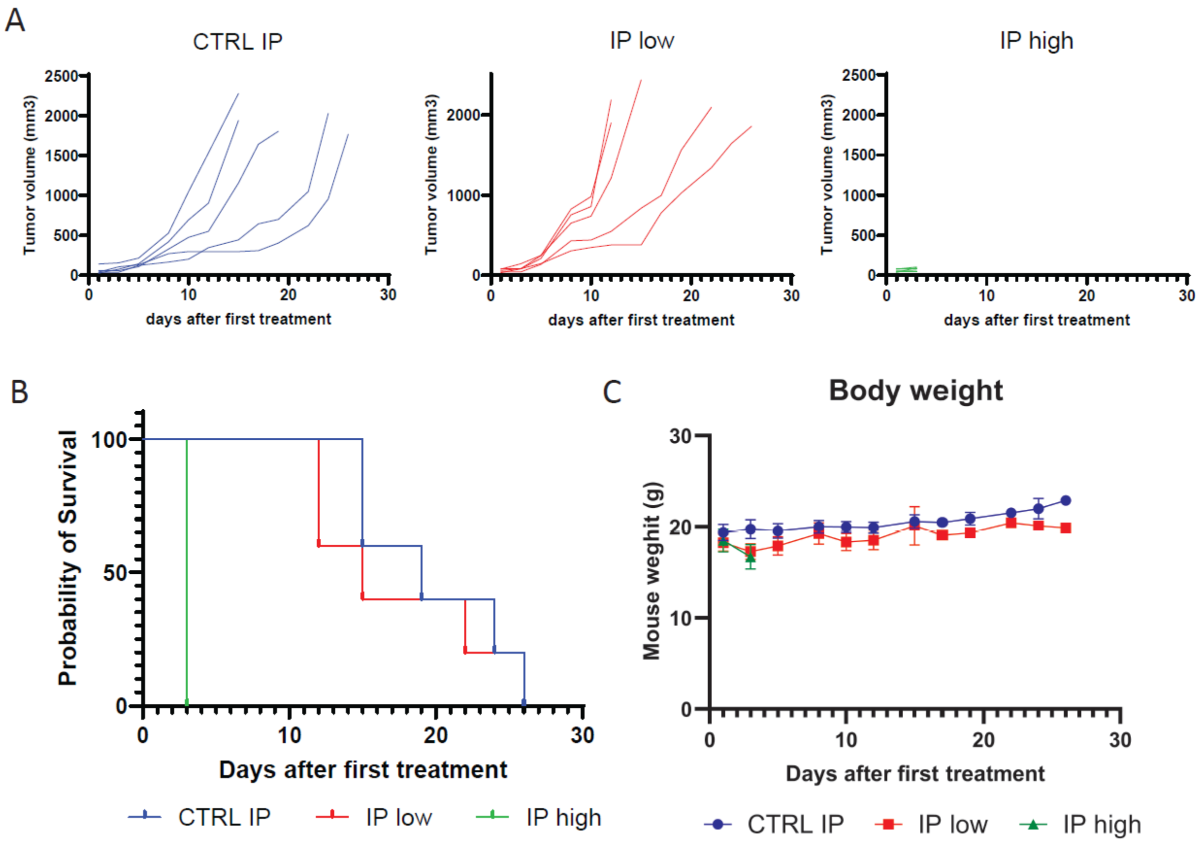

**Supplementary Figure 6. Assessment of KAT/3BP anti-lymphoma activity in PO and IT administration as single and in combination.** A) Tumor volume mean with positive and negative SEM for each group. B) Body weight mean with standard deviation for each group. Figures up to the end of the experiment (day 44).

A

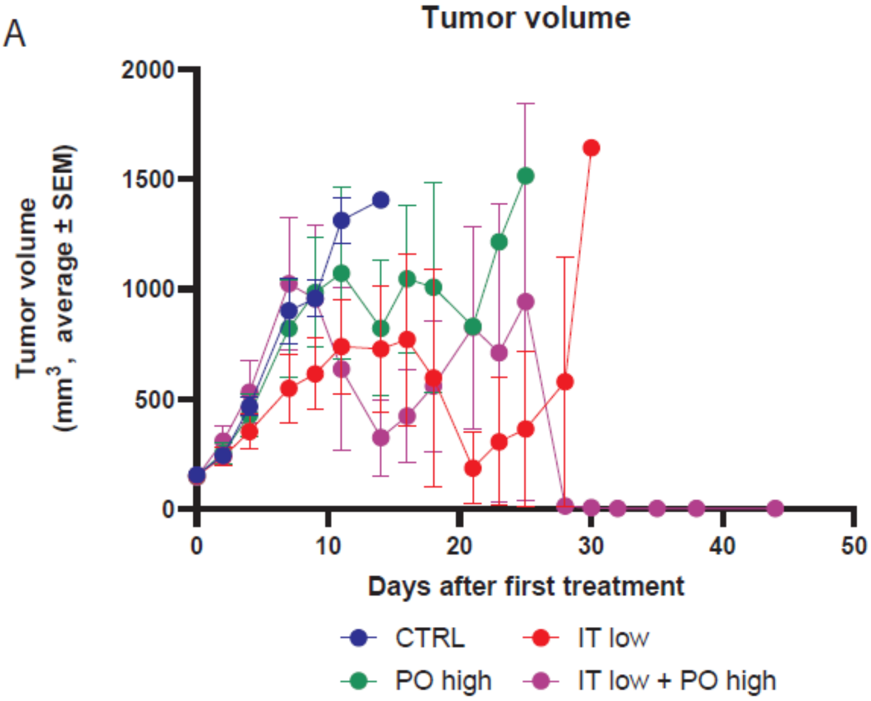


B

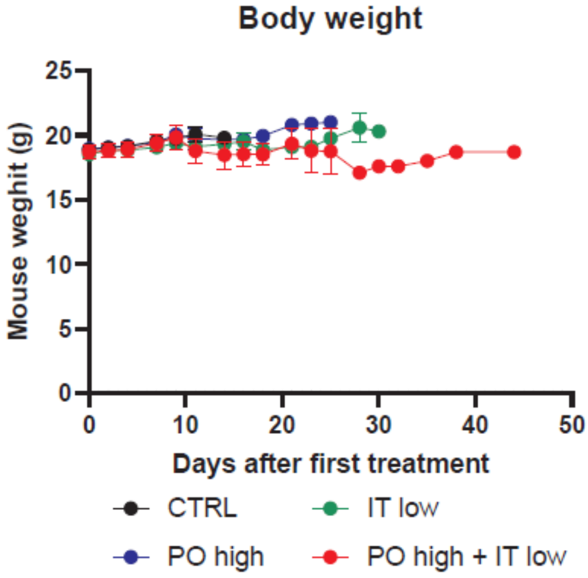

**Supplementary Table 1. Compounds combined with KAT/3BP with their respective target and range of concentrations used.** KAT/3BP was used at 30 μM 1:2 dilution. *, rituximab, cyclophosphamide, doxorubicin, vincristine, prednisone; ** in ABC-DLBCL, due to the drug clinical use; *** in GCB-DLBCL, due to the drug clinical use, and in the murine cell.

| Agent | Mechanism of action | Concentration range |
|---|---|---|
| Bendamustine | Chemotherapy | 20 μM (1:2 dilution) |
| R-CHOP * | Chemotherapy plus anti-CD20 monoclonal antibody | R, 20 μg/ml; CHOP, 2 μg/ml (1:2 dilution) |
| Venetoclax | BCL2 inhibitor | 10 μM (1:4 dilution) |
| Ibrutinib ** | BTK-inhibitor | 20 nM TMD8; 20 μM 1:2 U2932 (1:2 dilution) |
| Lenalidomide ** | Immunomodulatory | 20 μM (1:2 dilution) |
| Copanlisib | pan class I PI3K inhibitor (PI3Kδ/α) | 10 μM (1:4 dilution) |
| Tazemetostat *** | EZH2 inhibitor | 25 μM (1:2 dilution |
| Vorinostat | HDAC inhibitor | 10 μM (1:3 dilution) |

**Supplementary Table 2. Overall experimental design to assess the efficacy of KAT/3BP via various delivery routes (oral, IT, and IP) in tumor-bearing syngeneic mice.**

| Group n. | Total number of mice | compound | treatment | schedule | route |
|---|---|---|---|---|---|
| 1 | 5 | Vehicle (SFB) | / | 4 on 3 off, 4 weeks, 1 W recovery | PO |
| 2 | 5 | Vehicle (SFB) | / | 4 on 3 off, 4 weeks, 1 W recovery | IT |
| 3 | 5 | Vehicle (SFB) | / | 4 on 3 off, 4 weeks, 1 W recovery | IP |
| 4 | 5 | KAT/3BP | 2.5 mg/kg | 4 on 3 off, 4 weeks, 1 W recovery | PO (low) |
| 5 | 5 | KAT/3BP | 10 mg/kg | 4 on 3 off, 4 weeks, 1 W recovery | PO (high) |
| 6 | 5 | KAT/3BP | 0.5 mM | 4 on 3 off, 4 weeks, 1 W recovery | IT (low) |
| 7 | 5 | KAT/3BP | 2 mM | 4 on 3 off, 4 weeks, 1 W recovery | IT (high) |
| 8 | 5 | KAT/3BP | 2.5 mg/kg + 0.5 mM | 4 on 3 off, 4 weeks, 1 W recovery | PO (low) + IT (low) |
| 9 | 5 | KAT/3BP | 10 mg/kg + 2 mM | 4 on 3 off, 4 weeks, 1 W recovery | PO (high) + IT (high) |
| 10 | 5 | KAT/3BP | 2.5 mg/kg | 4 on 3 off, 4 weeks, 1 W recovery | IP (low) |
| 11 | 5 | KAT/3BP | 10 mg/kg | 4 on 3 off, 4 weeks, 1 W recovery | IP (high) |

**Supplementary Table 3. Overall experimental design to assess the efficacy of KAT/3BP via various delivery routes in combination (PO, IT, combination) in tumor-bearing syngeneic mice.**

| Group n. | Total number of mice | compound | treatment | schedule | route |
|---|---|---|---|---|---|
| 1 | 8 | Vehicle (SFB) | / | 4 on 3 off, 4 weeks, 1 W recovery | PO+IT |
| 2 | 8 | KAT/3BP | PO high 10.0 mg/kg | 4 on 3 off, 4 weeks, 1 W recovery | PO high |
| 3 | 8 | KAT/3BP | IT low 0.5 mM | 4 on 3 off, 4 weeks, 1 W recovery | IT low |
| 4 | 8 | KAT/3BP | PO high + IT low | 4 on 3 off, 4 weeks, 1 W recovery | PO+IT |
